# Dose-Dependent Effects of Transcranial Alternating Current Stimulation on Spike Timing in Awake Nonhuman Primates

**DOI:** 10.1101/696344

**Authors:** Luke Johnson, Ivan Alekseichuk, Jordan Krieg, Alex Doyle, Ying Yu, Jerrold Vitek, Matthew Johnson, Alexander Opitz

**Affiliations:** Department of Neurology, University of Minnesota, Minneapolis, MN, USA; Department of Biomedical Engineering, University of Minnesota, Minneapolis, MN, USA; Department of Neuroscience, University of Minnesota, Minneapolis, MN, USA

**Author notes:** Correspondence (Alexander Opitz).

## Abstract

Weak extracellular electric fields can influence spike timing in neural networks. Approaches to impose such fields on the brain in a noninvasive manner have high potential for novel treatments of neurological and psychiatric disorders. One of these methods, transcranial alternating current stimulation (TACS), is hypothesized to affect spike timing and cause neural entrainment. However, the conditions under which these effects occur in-vivo are unknown. Here, we show that TACS modulates spike timing in awake nonhuman primates (NHPs) in a dose-dependent fashion. Recording single-unit activity from pre-and post-central gyrus regions in NHPs during TACS, we found that a larger population of neurons became entrained to the stimulation waveform for higher stimulation intensities. Performing a cluster analysis of changes in interspike intervals, we identified two main types of neural responses to TACS – increased burstiness and phase entrainment. Our results demonstrate the ability of TACS to affect spike-timing in the awake primate brain and identify fundamental neural mechanisms. Concurrent electric field recordings demonstrate that spike-timing changes occur with stimulation intensities readily achievable in humans. These results suggest that novel TACS protocols tailored to ongoing brain activity may be a potent tool to normalize spike-timing in maladaptive brain networks and neurological disease.

## INTRODUCTION

It is increasingly recognized that local field potentials (LFPs) are not only a mere byproduct of synchronized neural activity but have an important function of controlling neural excitability^1,2^. This phenomenon, called ephaptic coupling, describes the coordinating influence of local electric fields or brain oscillations on neural spiking activity^3,4^. Spike timing is a key mechanism for neural encoding, communication in brain networks, and plastic changes^5,6^. Abnormal brain oscillations underlie a wide range of neurological and psychiatric disorders^7–9^ and tools to normalize pathological brain function have high potential for clinical applications.

It is possible to impose weak electric fields on the human brain in an intensity range comparable to endogenous fields using noninvasive technologies. One prominent method – transcranial alternating current stimulation (TACS), applies weak oscillating electric currents through electrodes attached to the scalp^10^. Invasive intracranial measurements have demonstrated that intensities typically employed in human studies (up to 2 mA^11^) can achieve electric field strengths reaching up to 1 mV/mm^12,13^ in the human brain. A long history of *in-vitro* studies and *in-vivo* measurements in rodent models^14–22^ has established that these weak electric fields are sufficient to induce acute physiological effects. *In-vitro* preparations (i.e. isolated cells or slices), however, lack crucial system-level properties. Rodent *in-vivo* studies are limited due to the use of anesthesia which affects calcium channels and neural dynamics. Furthermore, rodents have different brain anatomy and cytoarchitecture which can lead to a different affinity to external TACS electric fields^23,24^. Thus, it is not clear how these findings translate to *in-vivo* physiological effects in awake humans or nonhuman primates, where endogenous ongoing activity in complex oscillatory networks might either amplify or suppress the effects of weak externally applied electric fields.

Recent studies about the mechanisms of TACS are conflicting. One study using invasive electrophysiological measurements in surgical epilepsy patients did not find entrainment effects of TACS^25^, however only an indirect measure was used. Further, studies based on measurements in anesthetized rats claimed that TACS intensities are too weak^26^ or physiological responses arise from secondary stimulation effects of the peripheral nervous system^27^. On the other hand, a recent study performing hippocampal recordings in awake nonhuman primates^28^ showed that TACS can directly entrain single-neuron activity in a frequency-specific manner. However, TACS electric fields using the same stimulation montage are typically larger in nonhuman primates than in humans^12,23^ and findings in deep brain regions could be mediated by cortical effects experiencing higher electric fields. Thus, an investigation of direct stimulation effects across a range of electric field intensities in neocortex within or close to a human applicable range is crucial to establish TACS as a meaningful tool to affect neural activity.

Here, we conduct dose-response measurements of single-unit activity in the pre-and postcentral gyrus of awake nonhuman primates during TACS. Our approach overcomes several limitations of previous studies: 1) We perform recordings in awake nonhuman primates. It is well known that anesthesia affects neural excitability which can strongly influence the response to TACS. Further, a nonhuman primate model is more closely related to the human brain where gyrification is important for brain stimulation effects^29^. 2) Single-unit recordings are a direct electrophysiological measure of neural activity which can be recovered in the presence of a stimulation artifact^22,28^. This is in contrast to previous studies measuring local field potentials or scalp EEG recordings in humans, which are heavily affected by complex artifacts^25,30^. 3) Simultaneous electric field recordings control for the effective stimulation dose^31^ and allow for direct comparison to human applicable stimulation parameters.

Our findings demonstrate the effectiveness of TACS to affect neural spike-timing and entrain spikes at the applied stimulation frequency in a dose dependent fashion. With increasing electric field intensity, more neurons respond to stimulation. Finally, by clustering interspike interval changes across neurons, we identify two independent responses to TACS: increased burstiness and phase entrainment. The findings are a first mechanistic study of dose-response relationships of immediate effects of TACS in awake primates and provide important insight into the neurophysiological mechanisms of TACS.

## METHODS

### Surgical Procedures

All procedures were approved by the Institutional Animal Care and Use Committee of the University of Minnesota and complied with the United States Public Health Service policy on the humane care and use of laboratory animals. Two adult female nonhuman primates (*macaca mulatta*) were used in this study (Subject A: female, 10 kg, 18 years; Subject B: female, 12 kg, 21 years). The animals were implanted with a head post and cephalic chamber positioned midline over motor cortical areas. In Subject A, all implant materials were made of MRI-compatible, non-conductive PEEK and ceramic except for a single titanium bone screw; in Subject B the implant materials including headpost, chambers, and bone screws were made of titanium. After surgery in both animals, a 96-channel microdrive with independently moveable tungsten microelectrodes was fixed to the cephalic chamber (Gray Matter Research). Confirmation of the chamber placement and electrode positions was done by coregistering preoperative 7-Tesla T1 MRI and postoperative CT scans using the Cicerone software package^32^.

### Electrophysiological recordings and transcranial current stimulation

Neurophysiological data were collected using a TDT workstation (Tucker Davis Technologies) operating at ~25-kHz sampling rate. A large dynamic range (±500 mV) and 28-bit resolution allowed recording of electrophysiological data during TACS, fully capturing both neural data and stimulation artifacts without data loss (e.g. due to amplifier saturation). In Subject A, the common reference for the electrophysiological recordings was a titanium bone screw positioned anterior to the recording chamber; in Subject B, the common reference was the cephalic chamber. All data were collected during an awake, resting state while the animal was seated in a primate chair that kept the head facing forward. TACS was applied through two round stimulation electrodes (3.14 cm2, Ag/AgCl with conductive gel [SignaGel]) attached over the skin of the left and right temple (Fig. 1A). This electrode montage was chosen to minimize effects of the implant and skull defects on the TACS electric fields^33^ by applying currents in an orthogonal direction to the recordings^13^. In addition, performing concurrent electric field recordings (see below) allowed us to directly measure and control the biophysical stimulation condition. This ensured that results would be comparable to stimulation parameters commonly used in human TACS studies.

**Figure 1:**
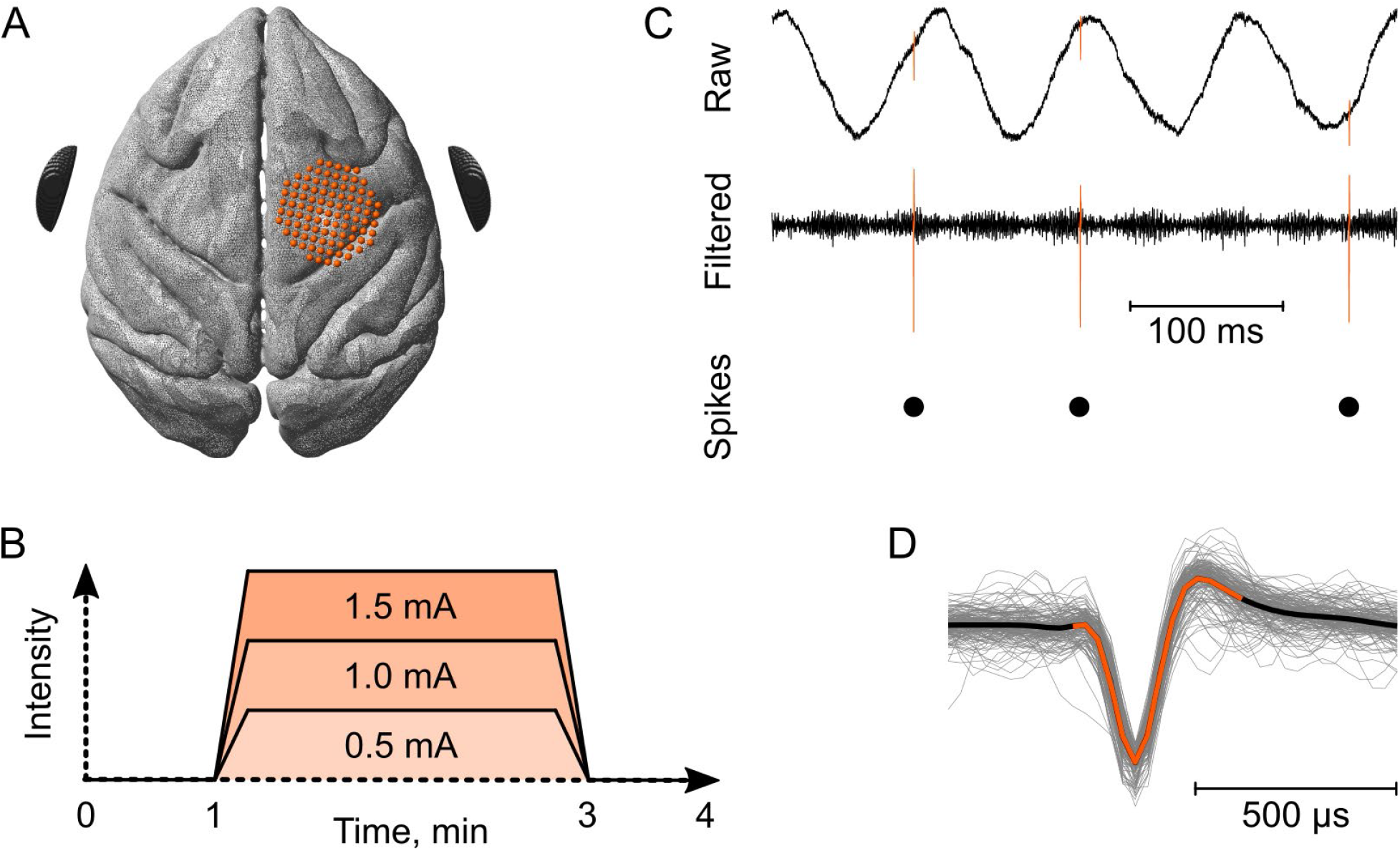
Experimental outline and data processing. **A)** A 96-channel microelectrode microdrive (maroon dots indicate different contacts at their original cortical location) was used to record from motor cortical regions (pre-motor and primary motor cortex). TACS was applied through two round stimulation electrodes (black) attached on the scalp over the left and right temples. **B)** Stimulation time course of different conditions. TACS was applied at 0.5 mA, 1.0 mA or 1.5 mA for two minutes with resting periods before and after stimulation. **C)** Recorded raw data of neural activity (orange) in the context of the tACS artifact and other noise sources (black). Signal filtering suppressed the tACS artifact allowing spike extraction, sorting, and identifying time stamps for spike times with respect to the stimulation phase. **D)** Overlaid spike waveforms before, during, and after TACS (from one exemplary neuron) demonstrating the consistency of spike waveforms in the presence of the stimulation artifacts.

At the beginning of the experiments, we recorded a rest condition without stimulation (0 mA) for several minutes. Then, TACS was applied at a frequency of 10 Hz for 2 min with 5s ramp up and down of the current amplitudes. Stimulation intensities applied were 0.5 mA, 1.0 mA and 1.5 mA (peak-to-baseline, Fig. 1B). Before and after each TACS run, resting condition was recorded for several minutes. In addition, a peripheral control stimulation was conducted where TACS was applied at 10 Hz and 1mA intensity through two electrodes attached over the shaved right upper arm in subject B.

### Data Analysis

One key issue to evaluate neurophysiological TACS effects measured with electrophysiology is the recovery of neural activity in the presence of a stimulation artifact. TACS induces a narrow band sinusoidal signal in the range of several mV^12,34^ at the applied stimulation frequency. Due to the presence of harmonics and interactions with other physiological signals (heartbeat, breathing)^30,35,36^, a full recovery of LFP data in the spectral range of interest is challenging. Spikes are in a frequency range much higher than the TACS signal thus filtering can suppress most of the stimulation artifact (see Fig. 1C for graphical outline of data processing). Even more important, spikes have a characteristic waveform which can identify them in the presence of other superimposed signals (Fig. 1D). Due to these considerations, we focused our analysis on spiking activity during TACS.

#### Spike Sorting

Within the 96-channel microdrives, neuronal spike recordings were collected from 15 (Subject A) and 11 (Subject B) microelectrodes that were inserted in pre-and post-central gyrus regions. These wide-band recordings were then analyzed offline using custom software developed in MATLAB (Mathworks) and Offline Sorter (Plexon). Raw signals were bandpass filtered between 0.3-10 kHz in Subject A and 0.3-5 kHz in Subject B, and single units were isolated and sorted using principal component and template-based methods in Offline Sorter. In subject B, TACS artifacts were larger than in subject A (likely due to the difference in recording ground). As such, a comb filter was also used to remove artifact harmonics that bled into the neural spike recording spectrum (1-5 kHz). We identified 38 neurons (n = 17 for animal A and n = 21 for animal B), of which four where excluded due to very sparse firing (< 0.25 Hz) not resulting in a sufficient number of spikes in the studied time period (final n = 15 for animal A and n = 17 for animal B).

#### Electric field recording

All electrodes within the microdrive were used for estimating the electric fields during TACS. The recorded data was bandpass filtered in a narrowband around the applied stimulation frequency (10 Hz ± 1 Hz) using a forward-reverse, zero phase, second order Butterworth IIR filter to derive the TACS voltage signal^12^. This resulted in a three-dimensional voltage distribution at the location of the recording electrodes. Electric fields in the recording area were then estimated by linearly interpolating the recorded voltages in a regular 3D grid and calculating the numerical derivative across grid points (Fig. 2).

**Figure 2:**
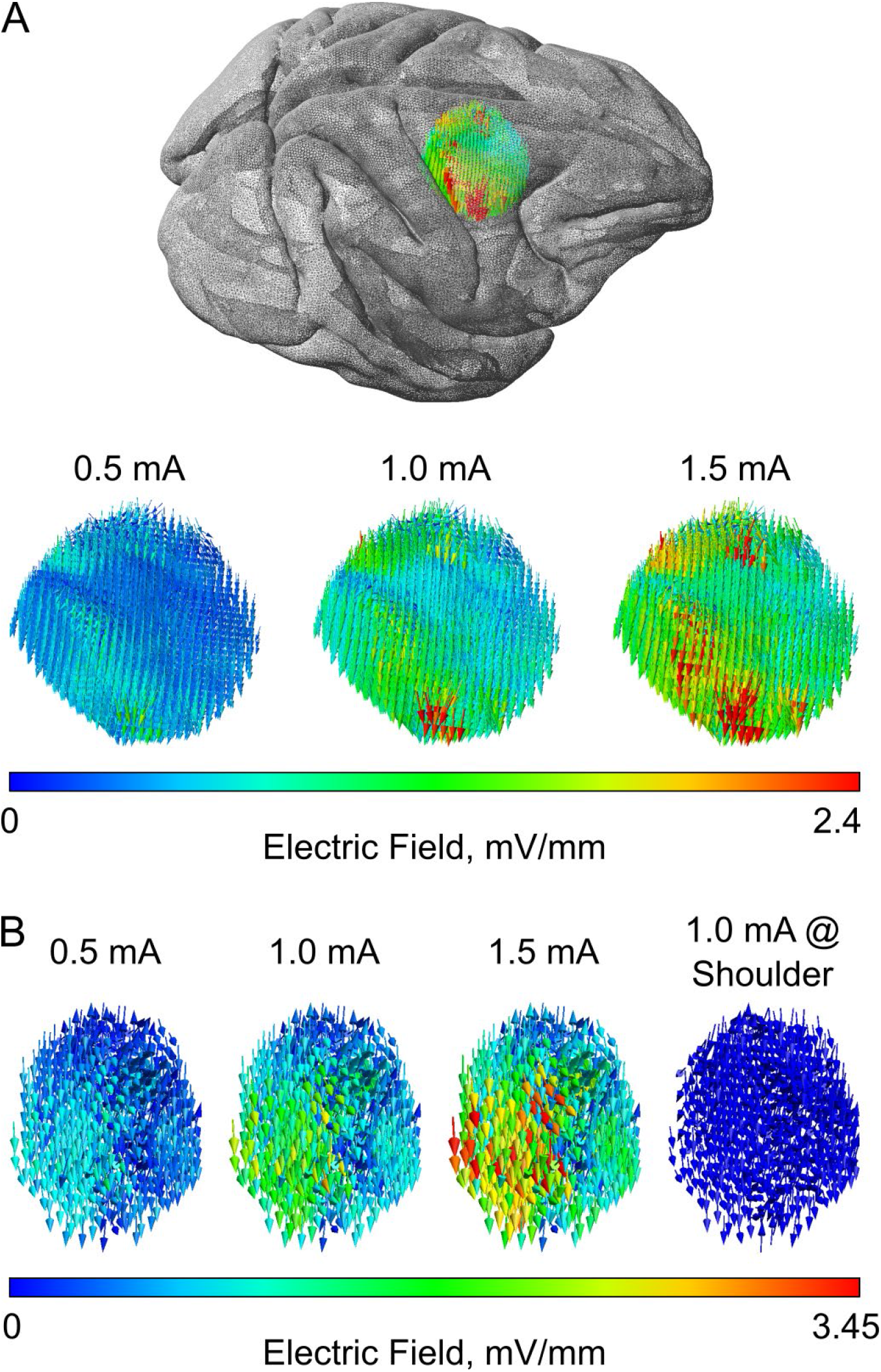
Recorded electric field during TACS in the neocortex. Electric field strength is color-coded with red indicating maximum electric field strength. **A)** Electric field distributions in Subject A on the brain surface for three different intensities (0.5 mA, 1.0 mA, 1.5 mA). The distributions have the same orientation and relative spatial relationships and scale linearly with intensity. **B)** Electric field distributions in Subject B for three different intensities and for the shoulder control stimulation.

#### Dose-response of spike entrainment to stimulation phase

In a first analysis, we evaluated dose-response effects of spike entrainment to TACS. For this, we determined the relative onset of spike times with respect to the TACS signal and computed peristimulus histograms. These show the number of spikes per time period of the stimulation. Deviation from uniformity indicates a preference for spiking relative to distinct stimulation phases. To compare the stimulation-induced changes to the resting ‘control’ condition, we determined spike times at rest (pre and post stimulation) relative to a virtual sinusoid with the same frequency as during TACS. In addition, we computed phase histograms indicating the preferred TACS phase during spiking (0-degree equals peak of stimulation). Further, we computed firing rates and phase lag values (PLV) pre, post, and during stimulation for all neurons. We identified neurons as responding to TACS based on a Rayleigh uniformity test (Rayleigh value > 4.6, p < 0.01) using the circular statistics toolbox^37^. These analyses were performed and compared across all three stimulation intensities (0.5 mA, 1.0 mA, 1.5 mA) and during the shoulder stimulation (1.0 mA). To test the statistical significance of changes in Rayleigh z values, PLVs, and firing rates, we implemented three generalized linear mixed effect models (glmm). Each outcome variable had 306 observations (34 neurons x 3 intensities x 3 conditions, which are the pre-, during, and post stimulation periods). In each model, the stimulation intensity (0.5, 1.0, or 1.5 mA) was the fixed effect factor, and the presence of stimulation (on or off), the subject id (subject A or B), and the membership of the neuron to an inter spike interval cluster (1^st^ or 2^nd^ cluster, see next paragraph) were the random effect factors to account for possible interindividual variability in the neural responses. The model parameters were estimated using a restricted maximum pseudo likelihood procedure with a log link function. All F values, degrees of freedom, and p values are reported in the main text.

#### Interspike intervals changes during TACS

While entrainment is one of the key postulated mechanisms for TACS^38^, it is conceivable that other changes in spike timing can occur due to stimulation. To study this, we computed interspike interval histograms during rest and during TACS. Interspike intervals (ISIs) indicate the time difference between two subsequent spikes (e.g. 0.1 s if spike n is followed by a spike n+1 with a 100 ms delay). ISI histograms plot the ISI of spike n against the ISI of spike n+1. ISI histograms can give insights into the temporal spiking patterns of neurons (e.g. indicate burstiness if spikes are followed by fast subsequent ones or spiking periodicity when spikes occur at similar time delays). We computed ISI histograms during TACS and resting state conditions and estimated their difference (TACS – rest). To identify independent classes of neural responses, we computed a 2d cross-correlation matrix from the ISI histograms, which can be treated as a weighted graph. Using Newman’s modularity (community detection)^39^ implemented in the Brain Connectivity Toolbox^40^ we identified two main classes of TACS responses. All ISI difference histograms were then averaged within each class to create a mean class histogram. The robustness of the mean class histograms, and thus of the community detection, was assessed using a cluster-based permutation test (12,500 permutations). For this we shuffled the cluster labels, computed dummy mean class histograms, and compared the critical values of the real mean class histograms with those of the dummy mean class histograms. We defined the critical value of the histogram as the maximum cluster size – number of adjacent non-zero elements in the histogram after binarizing with a 90% intensity threshold.

## RESULTS

### Electric field recordings

Measured electric fields showed a dominant left-right orientation (Fig. 2A) which is expected given the location of the two stimulation electrodes. Median electric field strengths at the locations of the identified neurons were found to be 0.38 mV/mm, 0.77 mV/mm, and 1.15 mV/mm in subject A (Fig. 2A) and 0.43 mV/mm, 0.86 mV/mm, and 1.33 mV/mm in subject B for 0.5 mA, 1.0 mA and 1.5 mA, respectively (Fig. 2B). These values are in good agreement with previous invasive measurements or modeling studies^12,23^. According to Ohm’s law, the electric field strength increases linearly with intensity while the relative spatial distribution remains identical (Supplementary Figure 1). Maximum electric field strength was found towards the lateral edge which could be explained through the proximity of the stimulation electrode and gray matter-cerebrospinal fluid transition transition^41^.

### TACS dose-response relationship on spike entrainment

We evaluated the alignment of spike times with respect to the stimulation phase (“entrainment”). We found that several recorded neurons did spike closer in time to the TACS peak (0-degree) and less during the trough (Fig. 3). During both the TACS peak and trough, the electric field strength is maximal but with opposite direction. Depending on the orientation of the neuron with respect to the electric field, this is consistent with the idea of maximizing de- or hyperpolarization along the cortical neuron^42^. With increasing stimulation intensity, spiking activity became increasingly phase-locked to the TACS peak times and suppressed during the TACS trough (Fig. 3A). This entrainment effect occurred immediately after stimulation onset and ceased after offset. Of note, our stimulation lasted only 2 min, which is likely not enough to induce lasting effects. While entrainment to TACS was pronounced in several neurons it occurred only in a subset of all recorded neurons (Fig. 4). Thus, on the group level mean phase lag values over all recorded neurons increased only slightly during TACS (Fig. 4A). Identifying neurons based on their degree of phase entrainment (p < 0.01 in the Rayleigh test), we found that TACS at 0.5 mA, 1.0 mA, and 1.5 mA entrained 8.9%, 17.6%, and 26.5% of all neurons (n = 34), respectively. The generalized linear mixed effect model indicates a significant effect of stimulation intensity on the Rayleigh z-value (F_1,304_ = 11.54, p = 0.0008). The statistical model of PLV dependency from the stimulation intensity is also significant (F_1,304_ = 7.13, p = 0.008). In the responsive neurons, the mean PLVs were increasingly enhanced with higher stimulation intensities (Fig. 4B; mean PLV = 0.11, 0.15, and 0.19, respectively). During the pre-and post-stimulation periods, PLVs remained at the same, nonsignificant level (mean PLV = 0.06). Firing rates showed no consistent changes (Fig. 4C-D; glmm F_1,304_ = 0.08, p = 0.78). No changes in the spiking activity were found in the shoulder control stimulation condition (Fig. S2; all Rayleigh p > 0.1, mean PLV = 0.04).

**Figure 3:**
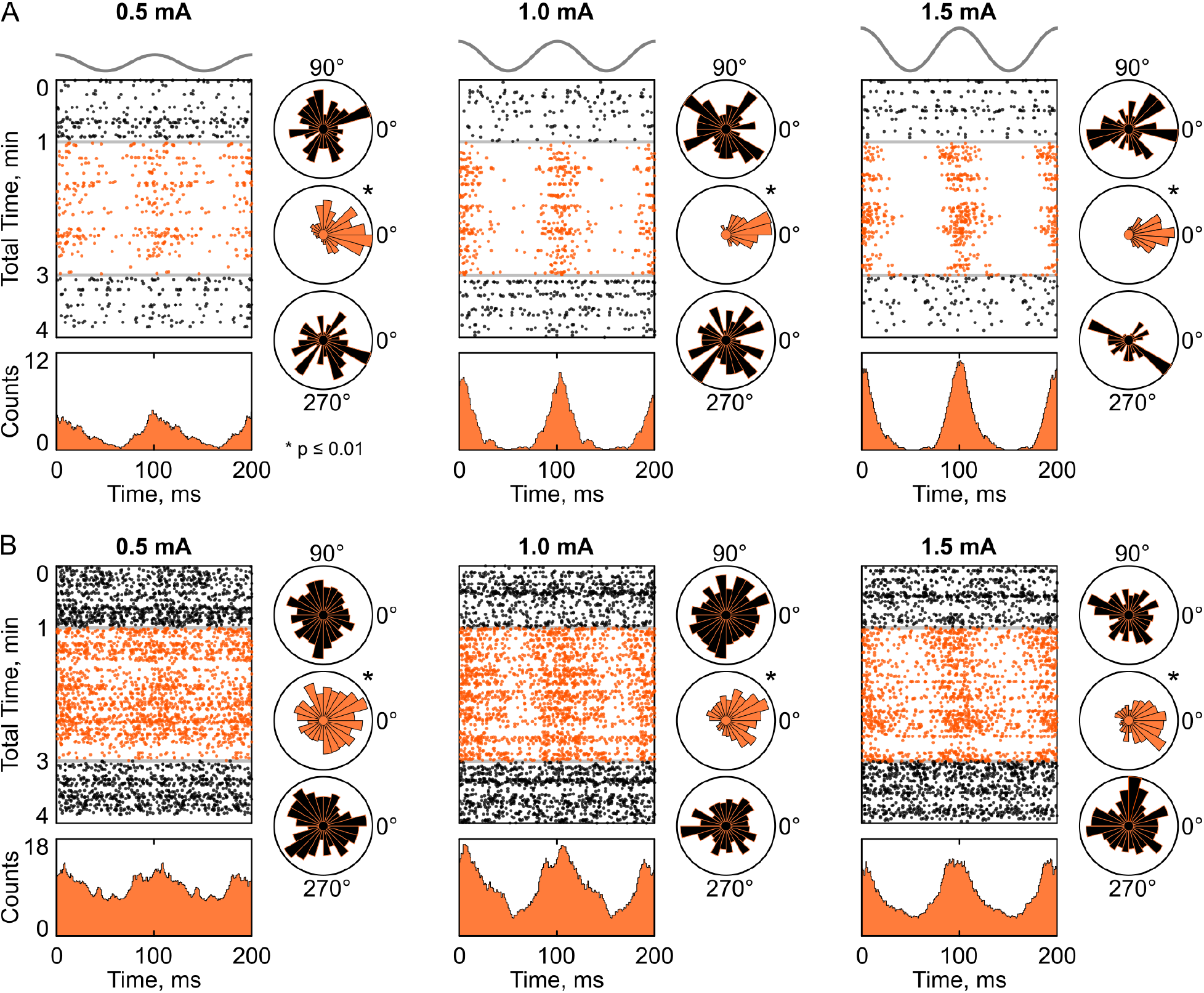
Dose-dependency of single-unit entrainment. **A)** Time-course of spikes (indicated as black dots for pre-/post-stimulation periods, and as orange dots – during stimulation) over two stimulation cycles (x-axis) over total recording time before, during, and after TACS (y-axis). The data is shown for one exemplary neuron for three intensities (0.5 mA, 1mA, 1.5mA). Pre-stimulation trials and post-stimulation trials (black) are arranged to a virtual stimulation signal with the same frequency (10 Hz) as during stimulation (orange). With stimulation onset spike times cluster more towards the peak of the stimulation cycle (corresponding electric field shown in Fig. 2) compared to a more uniform distribution during rest. Below, peristimulus histograms (smoothed with a 10 ms Gaussian kernel) of spike times during stimulation (TACS waveform is shown with gray above the figure) indicate a non-uniform distribution of spike onsets. On the right side, polar plots of the phase during spike onset are shown. * indicate significant (p < 0.01) non-uniformity according to a Rayleigh test. **B)** Same analysis as presented in A but for another exemplary neuron with higher spiking rate.

**Figure 4:**
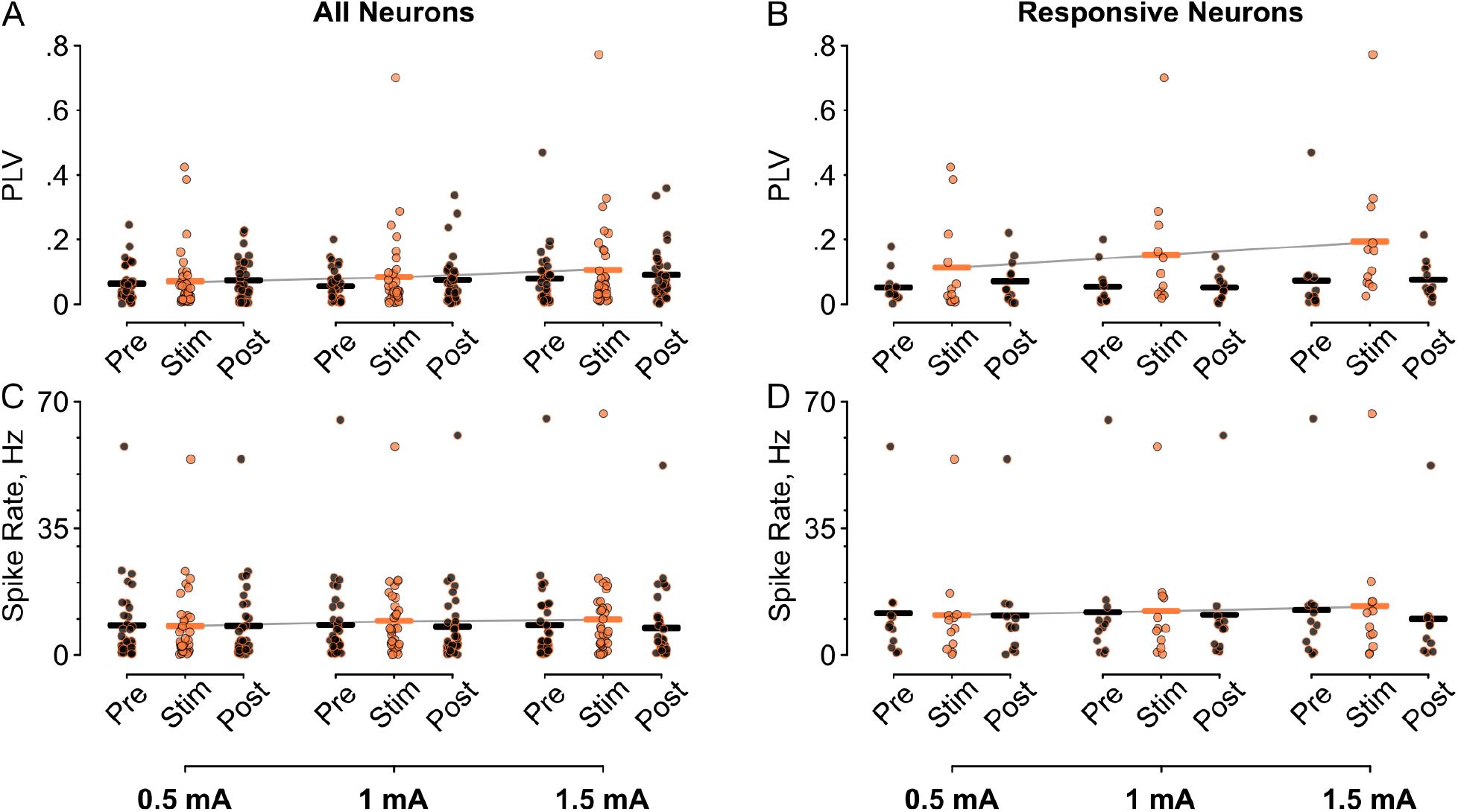
Phase-lag value and spike rate changes across all neurons. **A)** Phase-lag values across all recorded neurons (n = 34) are shown for all conditions (pre, during, and post stimulation) for three studied intensities (0.5 mA, 1.0 mA, 1.5 mA). Individual dots indicate values for each neuron and solid bars show the group mean. **B)** The increase in PLV during stimulation is driven by a subset of neurons responding to TACS with increased entrainment. PLV values during stimulation are enhanced during all intensities and increasingly so for higher intensities. **C)** Spiking rate during TACS and rest across all conditions are shown for all recorded neurons and **D)** for only responsive neurons. No consistent effects on spiking rate were observed during TACS. See Figure S2 for the same data during shoulder stimulation (control condition).

### Interspike intervals changes during TACS

Investigating a larger range of possible neural effects of TACS apart from entrainment, we calculated ISI histograms during TACS and at rest. We determined changes in spike timing induced by TACS by calculating the difference between ISI distributions in the on-and off-stimulation conditions. We identified principled types of neural responses to TACS by calculating Newman’s modularity^39^ based on the cross correlations of the ISI histograms. We identified two main classes of spike timing changes during TACS (Fig. 5): increased burstiness (Fig. 5A, see also Supplementary Fig. S3) and phase entrainment (Fig. 5B, see also Supplementary Fig. S4). With increasing intensities one class of neurons showed a shift towards shorter subsequent spike times during TACS compared to rest (n = 17 (9+8), 13 (5+8), and 18 (9+9) neurons (in subjects A + B) during 0.5 mA, 1.0 mA, and 1.5 mA, respectively). This is not accompanied, however, by a significant increase in overall firing rate. A second class of neurons showed entrainment to TACS with increased subsequent spike times at 100 ms and an accompanying decrease in faster spikes (n = 17 (6+11), 21 (10+11), and 16 (6+10) neurons, respectively). These two classes were found independently across all three stimulation intensities and were significantly non-random (cluster-based permutation test p < 0.01) at every intensity level. Analyzing the interspike intervals during shoulder stimulation, we found no meaningful classes of neural responses (p > 0.1).

**Figure 5:**
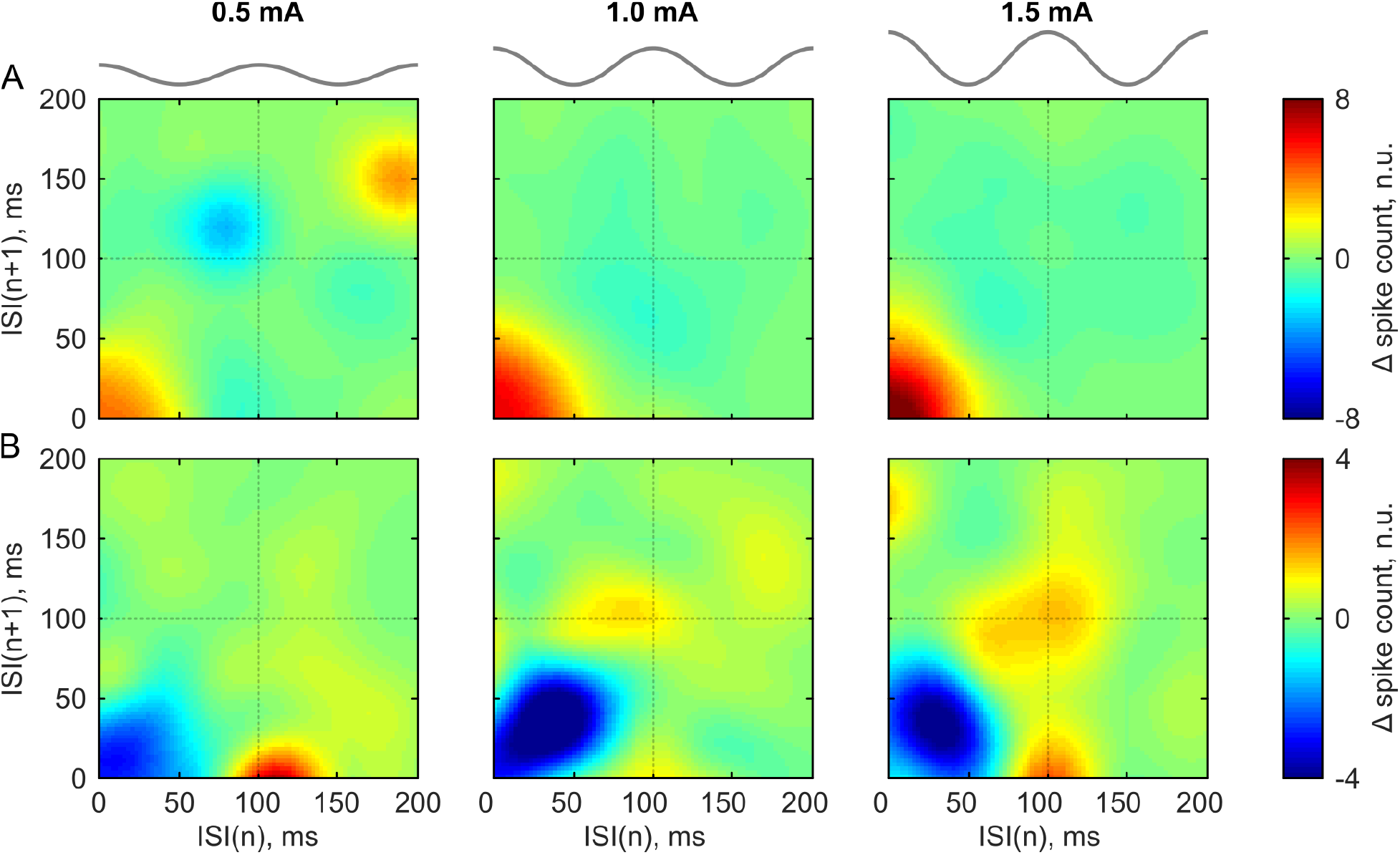
Two types of stimulation-induced interspike interval (ISI) changes (TACS minus rest). Each subplot shows a mean ISI histogram of neurons which belong to a given class according to Newman’s modularity analysis. All mean histograms are significantly non-random (cluster-based permutation test p < 0.01). Color coded are differences in spike counts (normalized units) of ISI histograms across all neurons falling into the class. **A)** Class 1. An increase in burstiness (spikes following each at low inter-stimulus times) during TACS is observed. This effect increases for higher stimulation intensities. **B)** Class 2. An increase in entrainment (enhanced ISIs at 100ms, based on 10 Hz waveform) is visible during TACS. This is accompanied with a decrease in fast successive spikes. See Figures S4 and S5 for the ISI histograms of individual neurons during TACS and rest.

## DISCUSSION

Conducting single-unit recordings in awake nonhuman primates we showed that TACS affects spike timing in a dose-dependent fashion. With increasing electric field strength, more neurons entrained to the stimulation frequency. Further, we identified increased burstiness as a second type of neural response to TACS. Importantly, concurrent electric field recordings demonstrated that these spike timing changes occurred in a field regime that are practicable in humans^23^.

One question of key importance for translational efforts to use TACS to affect brain rhythms (e.g. oscillopathies in schizophrenia^43^) is the intensity range at which physiological effects occur. If effective fields cannot be achieved in a safe and painless manner in humans, the therapeutic potential of TACS is severely hampered. Our electric field recordings indicate that entrainment effects can occur at field strengths < 0.5 mV/mm, which is achievable in humans for TACS intensities between 1-2 mA^12^. However, physiological effects were more pronounced for higher intensities, thus efforts to safely increase dosing have the potential for more consistent effects^38,44^.

Our data show that 10 to 20% of neurons respond to TACS with spike entrainment. Experiments *in-vitro*^14,42^ and modeling studies^45^ show that the responsiveness of neurons depend on their morphology and orientation to the applied electric field as well as their biophysical properties (e.g. ion channel dynamics). One possibility is that the entrained neurons were optimally aligned with respect to the electric field. Another possible reason is that their cell type-specific biophysical properties rendered them more susceptible to stimulation. Further, the increase of entrained neurons for higher field strengths fits well with the observation of a linear membrane depolarization for weak electric fields^46^. As a second neural response, we found increased burstiness of neurons. Burst firing of neurons has been observed in the presence of enhanced background activity in a low frequency range^47–49^. It is conceivable that in our recording TACS increased this background activity, thus resulting in the observed effect.

The time-course of our recordings indicates that entrainment occurred and ceased immediately with stimulation on-and offset, respectively. This does not support theories^50^ suggesting a buildup phase or stimulation echo. However, our protocol used short stimulation durations (2min) that were not intended to investigate the extent to which TACS elicits after-effects, such as due to spike-dependent plasticity^51,52^. Future studies systematically manipulating the TACS parameter range (frequency, electric field orientation, duration) will be important to identify the most effective stimulation parameters for immediate and lasting physiological effects. These studies will be facilitated by our increasing understanding of the underlying biophysics of TACS^12,53^ and sophisticated models to predict intracranial electric fields^41^.

It is unlikely that the observed physiological responses were due to peripheral nerve stimulation or other peripherally mediated effects. First, the alignment of spikes to the exact peak of the stimulation waveform (depolarization), as well as suppression during the trough (hyperpolarization) indicated a direct neural effect. For a peripheral pathway one would expect a time shift that reflects the conduction of the excitation through the neural system to the recording site in the neocortex, which should take at least several tens of milliseconds, similar to fast event-related potentials. Of note, our measured strength of entrainment, phase-lag values, was noticeably higher than in a preceding study in anesthetized rodents which found no direct neural effect of stimulation^27^. This suggests a largely unspecific effect and generally suppressed brain activity observed in this previous study. Second, the absence of entrainment in our shoulder stimulation control further corroborates a direct neural effect of TACS, which is also in line with another recent TACS study in awake NHPs^28^ performing additional control measurements using local skin anesthesia^54^. It is very likely that in an awake brain spike timing can be more easily affected compared to an anesthetized state used in previous studies^26,27^, thus possibly explaining differences found in effective stimulation doses across studies.

In conclusion, our results show that weak extracellular electric fields can affect spike-timing *in-vivo* in awake nonhuman primates in a dose dependent fashion. Enhancement or suppression of spiking activity during the peaks and troughs of stimulation suggest a neural membrane de- or hyperpolarization due to the applied weak extracellular fields. Entrainment effects were found in a subset of neurons that increased with stimulation intensity. Another robust neural response – increased burstiness, has not been considered so far in the TACS literature. This suggests the need for further systematic studies to examine TACS mechanisms in-vivo. Importantly, the found physiological responses occurred in an electric field regime applicable to human studies, thus providing an electrophysiological grounding for efforts to directly interact with human brain oscillations in a non-invasive and safe manner.

## ACKNOWLEDGEMENTS

Research reported in this publication was supported by the University of Minnesota’s MnDRIVE Initiative, in part by the National Institutes of Health (P50-NS098573, R01-NS058945), a predoctoral fellowship (NINDS, F31-NS108625) and Engdahl Philanthropic Donation. We thank Kelton Wilmerding and Shane Nebeck for assistance with the experimental preparations.

## AUTHOR CONTRIBUTIONS

L.J and A.O. designed the experiments; L.J, J.K, A.D and A.O. collected the data; L.J and M.J supervised the data collection; L.J., I.A., J.K., A.D., Y.Y. and A.O. processed MR and CT data and identified electrode locations; L.J, I.A. M.J. and A.O. analyzed the electrophysiological data and prepared the figures; M.J. and A.O. supervised the data analysis; L.J., I.A., M.J. and A.O. interpreted the results; A.O. wrote the paper with critical input from the co-authors; all authors reviewed the paper.

## SUPPLEMENTARY INFORMATION

**Figure S1:**
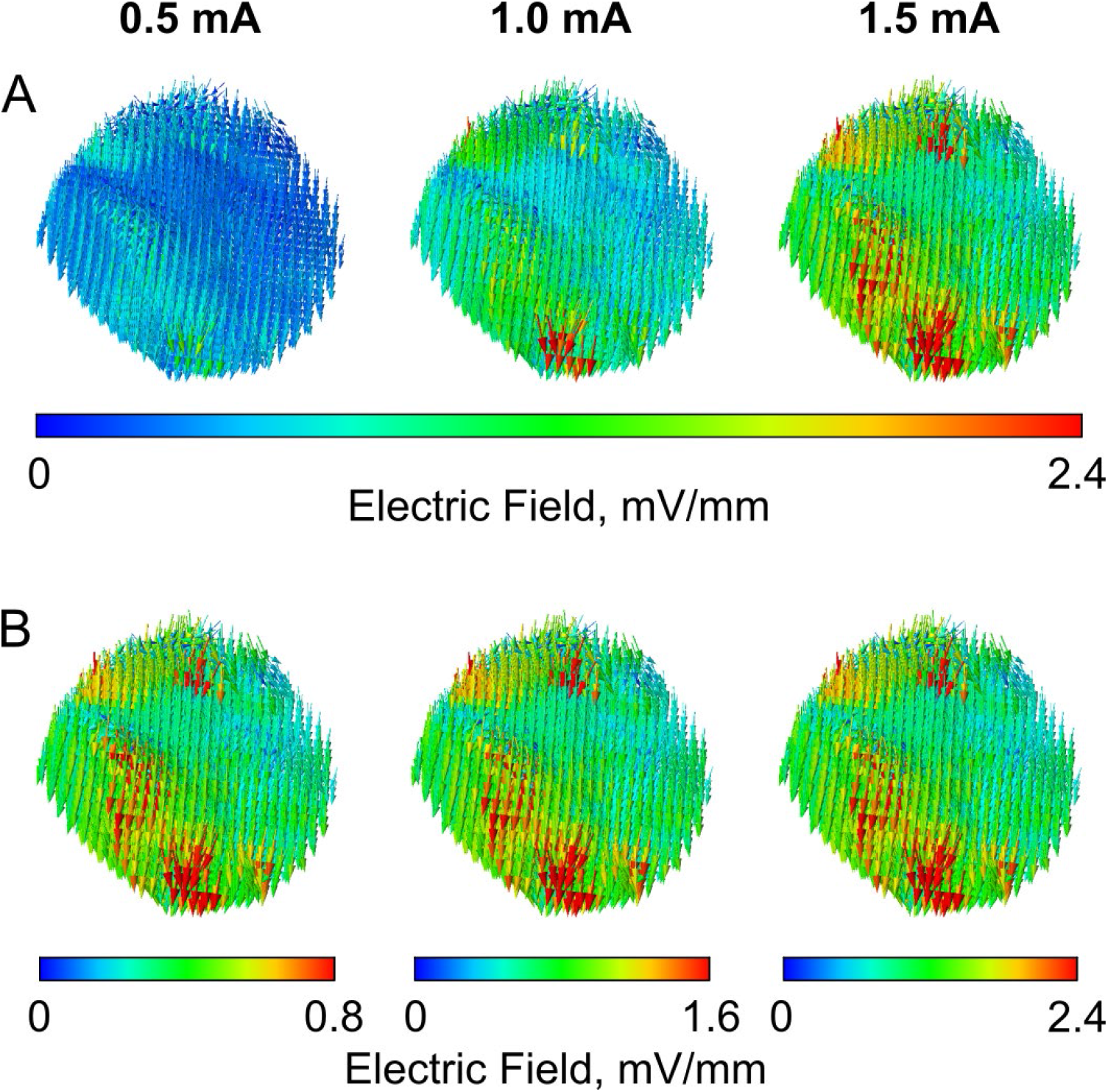
Recorded electric field during TACS in the motor cortex for Subject A on the same scale **(A)** and individual scales **(B)**. As expected, relative electric field distributions are identical across stimulation intensities. This figure corresponds to Figure 2 in the main paper.

**Figure S2:**
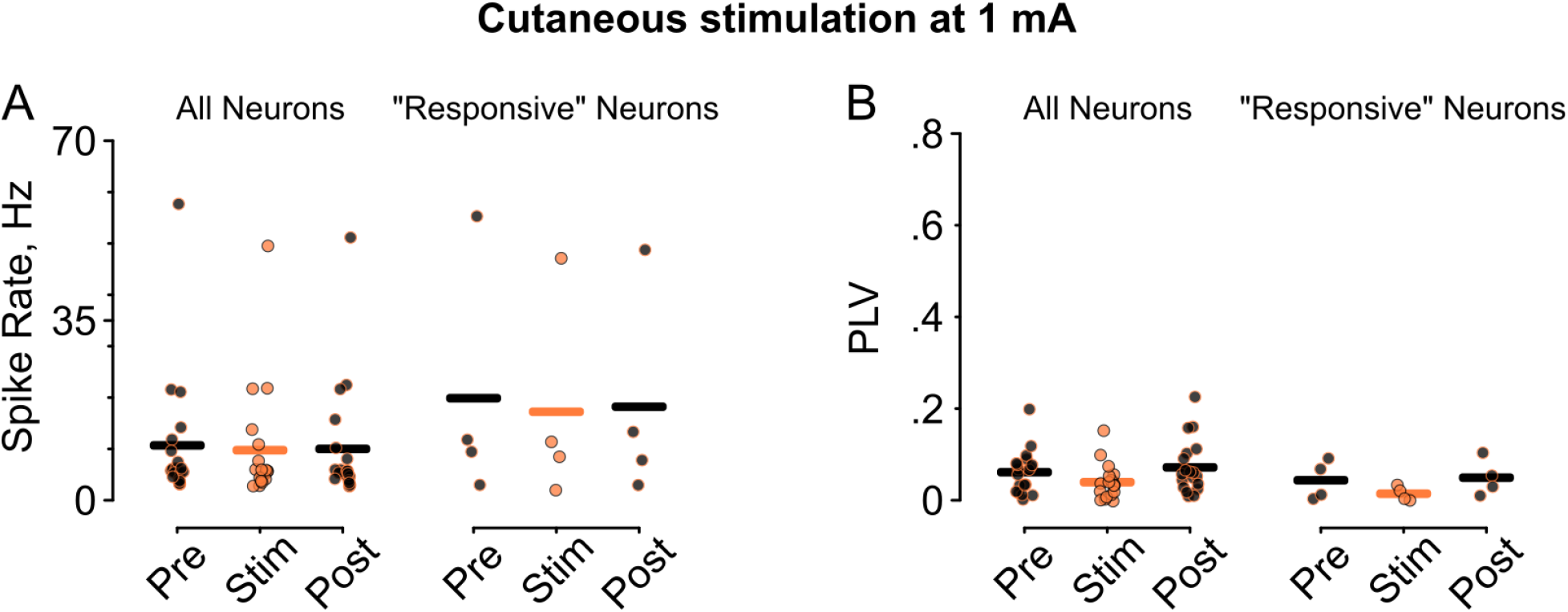
Spike rate changes **(A)** and phase-lag values **(B)** across all neurons (left) and neurons which responded to TACS (“responsive” neurons) during shoulder stimulation at an intensity of 1.0 mA. No changes in spike rate or changes in PLV (entrainment) during shoulder TACS was observed. This figure corresponds to Figure 4 in the main paper.

**Figure S3:**
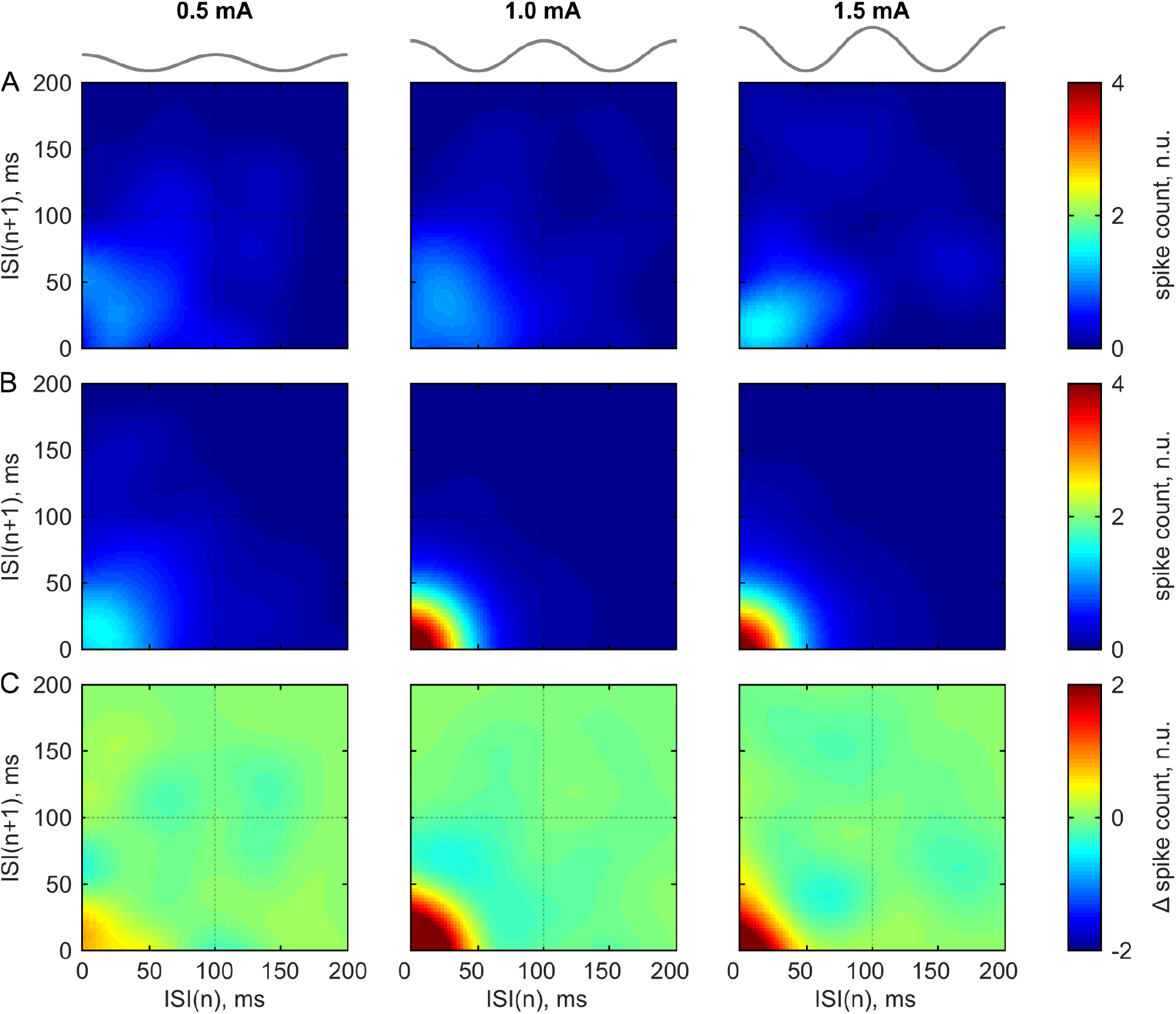
Interspike interval (ISI) histograms in one exemplary neuron during rest **(A)**, TACS **(B)**, and difference between TACS and rest **(C)**. Each column corresponds to one stimulation intensity (0.5 mA, 1.0 mA and 1.5 mA). Color coded are differences in spike counts (normalized units). This figure corresponds to a neuron in the class of cells (Class 1) represented in Figure 5A in the main paper.

**Figure S4:**
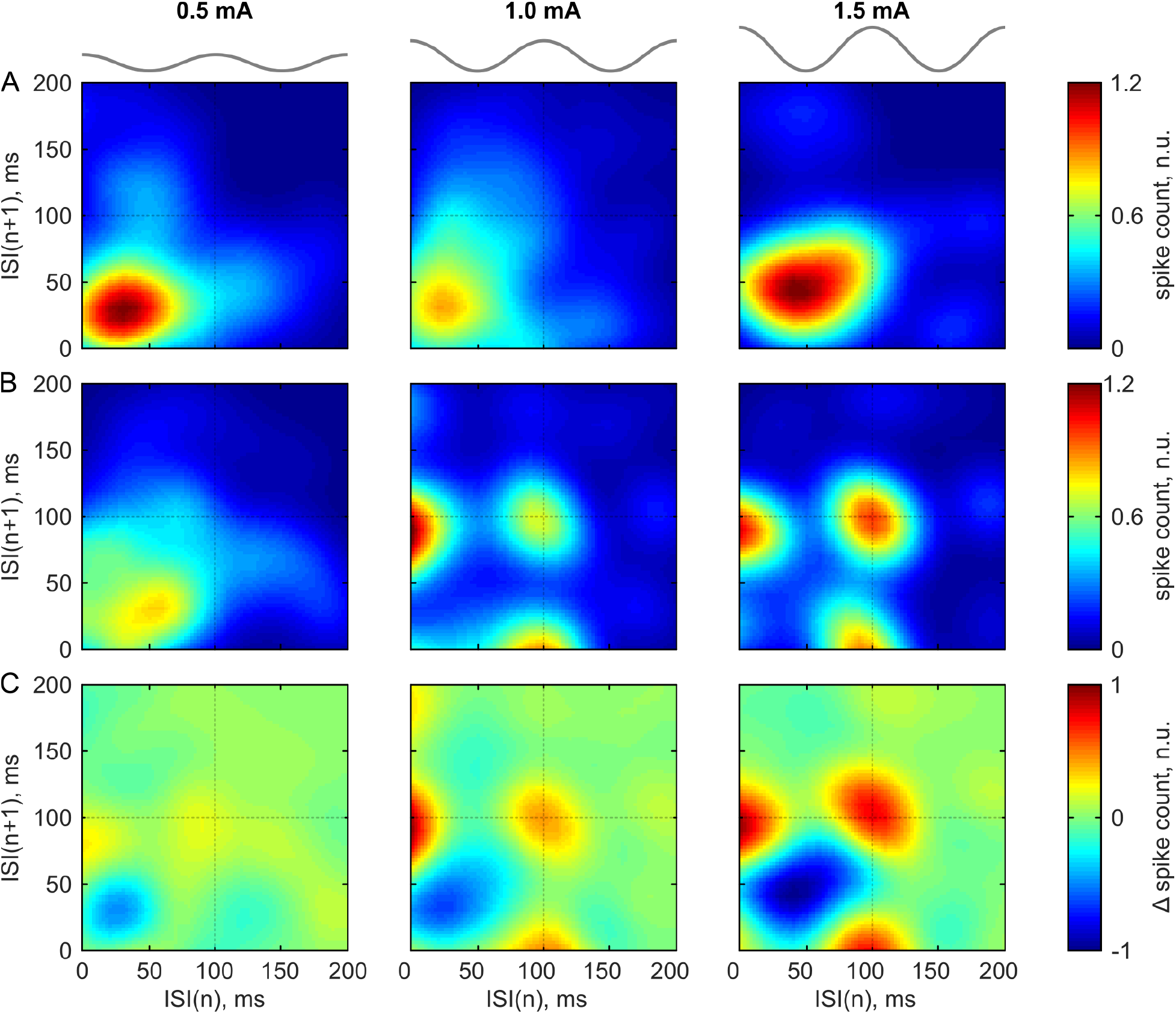
Interspike interval (ISI) histograms in one exemplary neuron during rest **(A)**, TACS **(B)**, and difference between TACS and rest **(C)**. Each column corresponds to one stimulation intensity (0.5 mA, 1.0 mA and 1.5 mA). Color coded are differences in spike counts (normalized units). This figure corresponds to a neuron in the class of cells (Class 2 represented in Figure 5B in the main paper.

